# The three *Plasmodium falciparum* Aurora-related kinases display distinct temporal and spatial associations with mitotic structures in asexual blood stage parasites and gametocytes

**DOI:** 10.1101/2024.05.27.596013

**Authors:** Matthias Wyss, Basil T. Thommen, Jacob Kofler, Eilidh Carrington, Nicolas M. B. Brancucci, Till S. Voss

**Author notes:** contributed equally to this work (Matthias Wyss, Basil T. Thommen).

## Abstract

Aurora kinases are crucial regulators of mitotic cell cycle progression in eukaryotes. The protozoan malaria parasite *Plasmodium falciparum* replicates via schizogony, a specialised mode of cell division characterized by consecutive asynchronous rounds of nuclear division by closed mitosis followed by a single cytokinesis event producing dozens of daughter cells. *P. falciparum* encodes three Aurora-related kinases (PfARKs) that have been reported essential for parasite proliferation, but their roles in regulating schizogony have not yet been explored in great detail. Here, we engineered transgenic parasite lines expressing GFP-tagged PfARK1-3 to provide a systematic analysis of their expression timing and subcellular localization throughout schizogony as well as in the non-dividing gametocyte stages, which are essential for malaria transmission. We demonstrate that all three PfARKs display distinct and highly specific and exclusive spatiotemporal associations with the mitotic machinery. In gametocytes, PfARK3 is undetectable and PfARK1 and PfARK2 show male-specific expression in late stage gametocytes, consistent with their requirement for endomitosis during male gametogenesis in the mosquito vector. Our combined data suggest that PfARK1 and PfARK2 have non-overlapping roles in centriolar plaque maturation, assembly of the mitotic spindle, kinetochore-spindle attachment and chromosome segregation, while PfARK3 seems to be exquisitely involved in daughter cell cytoskeleton assembly and cytokinesis. These important new insights provide a reliable foundation for future research aiming at the functional investigation of these divergent and possibly drug targetable Aurora-related kinases in mitotic cell division of *Plasmodium falciparum* and related apicomplexan parasites.

## Introduction

The life cycle of the protozoan parasite *Plasmodium falciparum*, the causative agent of the most severe form of malaria, includes multiple developmental stages in the *Anopheles* spp. mosquito vector and in humans as the intermediate host. During the symptomatic phase of the infection in the human bloodstream, merozoites invade red blood cells (RBCs) and develop within 48 hours from ring stage parasites into trophozoites and then schizonts that produce and release up to 30 merozoites determined to infect new RBCs (1). Repeated rounds of RBC invasion and intracellular replication result in the rapid increase of parasite loads, which is directly linked to malaria-related pathology (2). During each of these intra-erythrocytic replication cycles, a small subset of schizonts commit to gametocytogenesis and produce sexual ring stage progeny that differentiate within twelve days and through five morphologically distinct stages into mature transmission-competent female or male stage V gametocytes (3). Sexual commitment is initiated by the expression of the master transcription factor PfAP2-G (4, 5). The *pfap2-g* gene is generally kept in a silenced state by epigenetic factors including heterochromatin protein 1 (HP1) (6) and the histone deacetylase 2 (HDA2) (7). In response to changing environmental conditions, such as limiting levels of the host-derived lysophospholipid lysophosphatidylcholine (lysoPC) (8), however, the *pfap2-g* locus becomes activated in late trophozoites via a mechanism that involves the gametocyte development 1 (GDV1)-dependent eviction of HP1 (8, 9). Once expressed, PfAP2-G triggers a transcriptional response setting the course for sexual differentiation (10).

Intra-erythrocytic replication of *P. falciparum* blood stage parasites occurs via schizogony, a specialized mode of mitotic cell division that differs markedly from the process of binary fission through which mammalian cells divide (11–13). Schizogony begins when a trophozoite transitions from G1-phase into the first S-phase, proceeds with 4-5 consecutive asynchronous rounds of genome replication and nuclear division via closed mitosis, and concludes with a single cytokinesis event resulting in the segmentation of daughter merozoites from the multi-nucleated schizont (1, 12–16). Entry into and progression through schizogony critically depend on a centrosome-like structure referred to as the centriolar plaque (CP) that emerges in trophozoites (12, 15, 17–19). The CP represents a bipartite microtubule organising center (MTOC) that spans the nuclear envelope and consists of a cytoplasmic part (outer CP) and an intranuclear part (inner CP or intranuclear body) (13, 17, 18, 20–25). In trophozoites and schizonts, the centromeres of all 14 chromosomes assemble into a single perinuclear cluster that is inherently positioned in direct proximity to the CP (17, 22, 26, 27). At the late trophozoite stage, microtubules (MTs) extend from the intranuclear body into the nucleus to form a distinctive pre-mitotic structure called the hemispindle (13, 17, 18, 21, 28). During S-phase, the hemispindle retracts and the outer CP develops and separates into two branches, followed by duplication of the inner CP at the onset of mitosis (12, 18). During mitosis, the duplicated CPs nucleate a short bipolar mitotic spindle that connects to the kinetochores assembled at the centromeres of the sister chromatids (metaphase), which subsequently segregate to the opposite poles of the elongated nucleus now marked by an extended interpolar spindle (anaphase) (17, 18, 21, 27, 29, 30). At the end of the first mitotic cycle, nuclear fission produces two daughter nuclei, each equipped with a CP and ready to undergo a next round of genome replication and mitosis (17, 18, 21). The outer CP plays a crucial role in the biogenesis and faithful segregation of the apical complex, cellular organelles and cell division machinery during schizogony and segmentation (18, 19). Centrins, which belong to the few centrosomal components evolutionary conserved in malaria parasites, serve as reliable markers for visualization of the CP (31) and have recently been shown to localize exclusively to the outer CP (17, 18, 21, 32).

The CP is also present in gametocytes, which remain in G1-phase throughout the 12-day sexual differentiation process. In stage I gametocytes, the inner CP nucleates a long bundle of intranuclear MTs that spans the entire nucleus in stage II/III gametocytes and is degraded thereafter (22, 33). While the exact function of this non-mitotic hemispindle-like structure is still unknown, it appears to drive elongation of the gametocyte nucleus and participate in spatial genome organization by sequestering kinetochores (22). The outer CP nucleates the subpellicular MTs (SPMTs) that develop into an elaborate cytoskeletal network encasing the entire parasite in stage IV gametocytes before being disassembled again in mature stage V gametocytes (22, 23, 33, 34). During male gametogenesis, characterized by three rapid rounds of genome replication and endomitosis followed by karyokinesis of the octoploid nucleus and exflagellation of eight microgametes (35), the intranuclear body nucleates spindle MTs and the outer CP builds the basal bodies responsible for axoneme assembly (23, 36).

Progression through the mitotic cell cycle is controlled by several serine/threonine protein kinases collectively known as mitotic kinases, including the Aurora/Aurora-related kinase family. Aurora kinases are widely conserved in eukaryotes, with one member present in yeasts, two members in non-mammalian metazoans and three members in mammals (Aurora A, B and C) (37). Aurora A localizes to the centrosome and spindle poles and controls centrosome maturation and separation and bipolar spindle formation. Aurora B is a member of the chromosome passenger complex (CPC) (together with INCENP, Borealin, Survivin), localizes to the centromeres, kinetochores and spindle midzone and plays crucial roles in regulating chromosome condensation, kinetochore attachment, sister chromatid segregation and cytokinesis. Aurora C is closely related to Aurora B and specifically active in germ cells (37, 38). Based on sequence homologies among the catalytic kinase domains, three Aurora-related kinases (ARK1-3) have been identified in *Plasmodium* spp. and other apicomplexan parasites including *Toxoplasma gondii* (39, 40). Gene knockout screens performed in *P. falciparum* and *P. berghei*, a malaria parasite infecting rodents, indicated that all three parasite ARKs are likely essential for parasite proliferation (41–44). Importantly, targeted analyses demonstrated their localization to mitotic structures (36, 38–40), suggesting that ARKs are important regulators of cell division also in these divergent unicellular pathogens.

Reininger and colleagues demonstrated that PfARK1 is not expressed in ring stages and trophozoites but exclusively in schizonts, where PfARK1 expression was only observed in a subset of nuclei throughout schizogony. In these nuclei, PfARK1 localized to two closely paired foci flanking the short mitotic spindle, indicating that PfARK1 is associated with the spindle poles/intranuclear body specifically during the early phase of mitosis (39). Based on these observations, the authors speculated that PfARK1 may represent a functional homologue of mammalian Aurora A (38, 39). PfARK1 has also been reported as a target of the human Aurora kinase inhibitor hesperadin that blocks nuclear division in *P. falciparum* (45). *In vitro* selection for hesperadin-resistant parasites followed by whole-genome sequencing revealed resistance-conferring mutations in either PfARK1 or PfNEK1, a phylogenetically unrelated protein kinase (45). Whether these findings point at epistatic interactions between PfARK1 and PfNEK1 as suggested remains unclear (45), but they highlight these kinases as potential antimalarial drug targets (38, 45, 46). Furthermore, a recent preprint demonstrated that PbNEK1 also localizes to the CP and is essential for MTOC organization, formation of the mitotic spindle and kinetochore attachment during microgametogenesis in *P. berghei* (47). The localization and function of PfARK2 in *P. falciparum* has not yet been explored. Interestingly, however, *pfark2* transcription is upregulated in response to lysoPC depletion alongside with genes involved in sexual commitment and phospholipid metabolism, suggesting PfARK2 might be involved in one of these processes (8). In *P. berghei*, PbARK2 localization and function has recently been studied in detail during male gametogenesis (36). During this process, PbARK2 dynamically localizes to the spindle poles and laterally along the mitotic spindle in close juxtaposition to the attached kinetochores and the nuclear MT-binding protein PbEB1, a divergent homolog of the MT plus end-binding protein EB1 (36, 48). Transgenic *P. berghei* expressing reduced levels of PbARK2 showed no apparent defects in microgametogenesis and ookinete development, but oocyst formation and sporogony in the mosquito vector was severely impaired and completely blocked, respectively (36). PfARK3 was shown to be expressed in *P. falciparum* asexual blood stage parasites exclusively in multi-nucleated late stage schizonts, where it localized to single or paired foci or to slightly elongated structures in proximity to the nuclei (40).

Here, we generated conditional knockdown mutants to investigate the expression, subcellular localization and function of the three *P. falciparum* Aurora-related kinases PfARK1-3 in more detail. While inefficient knockdown of protein expression levels unfortunately prevented us from obtaining new insight into PfARK1-3 function, we were still able to monitor their expression and subcellular localization throughout intra-erythrocytic parasite development and gametocyte differentiation. Our results reveal distinct and dynamic expression and localization patterns in close association with mitotic structures for all three PfARKs, as well as male-specific expression of PfARK1 and PfARK2 in late stage gametocytes.

## Results

### Generation of inducible conditional PfARK knockdown cell lines

We designed a CRISPR/Cas9-based gene editing strategy to engineer *P. falciparum* mutants suitable to study the expression and subcellular localization of green fluorescent protein (GFP)-tagged PfARK1-3 at wild type expression levels, while at the same time facilitating their functional characterization using an inducible conditional protein knockdown approach (icKD). To this end, we combined the DiCre/loxP system for targeted sequence excision at the gene of interest (GOI) in response to rapamycin (RAPA) exposure (49–51) with the FKBP-derived destabilization domain (DD) system for conditional depletion of the protein of interest (POI) upon removal of the stabilizing ligand Shield-1 (52, 53). This approach is based on fusing to the 3’ end of the GOI a modular sequence tag that consists, in the following order, of (1) a loxP element inserted into the *sera2* intron (loxPint) (50); (2) the *gfp* coding sequence followed by a transcriptional terminator; (3) a second loxPint element; and (4) a sequence encoding a triple hemagglutinin tag fused to the DD domain (3xHA-DD) (Fig. S1). Using this setup, transgenic parasites can be used to study the temporal expression dynamics and subcellular localization patterns of the GFP-tagged POI under wild type expression levels. After RAPA-induced DiCre-mediated recombination of the loxP sites, parasites will express the POI-3xHA-DD fusion protein, allowing for functional studies by comparing parasites grown under protein stabilizing (+Shield-1) or knockdown (−Shield-1) conditions. Here, we introduced this modular sequence tag separately at each of the three *pfark* loci, resulting in the three transgenic cell lines NF54/ARK1-GFP-icKD, NF54/ARK2-GFP-icKD and NF54/ARK3-GFP-icKD (Fig. S1). As a parental strain, we used NF54/DiCre parasites (54) further modified to express mScarlet-tagged PfAP2-G allowing for the automated quantification of sexual commitment rates (SCRs) in live cells using high content imaging (55). Successful gene editing was confirmed for each of the three parasite lines by polymerase chain reaction (PCR) on parasite genomic DNA (gDNA) (Fig. S1).

### PfARK expression and localization in asexual blood stage parasites

Using live cell fluorescence microscopy, PfARK1-GFP expression was first detected in single-nucleated parasites at the late trophozoite/early schizont stage, where it localized to a single or two closely paired spots at the nuclear periphery (Fig. 1A). These distinct perinuclear PfARK1-GFP foci were observed throughout schizogony but only in a subset of nuclei in individual schizonts. In segmented schizonts, PfARK1-GFP signals were undetectable (Fig. 1A). Co-staining of PfARK1-GFP and Centrin in indirect immunofluorescence assays (IFA) revealed that PfARK1-GFP is closely associated but does not co-localize with a subset of the outer CPs and most frequently localizes in between duplicated CPs (Fig. 1B-C). Results from triple labeling IFAs using antibodies against α/β- tubulin showed that PfARK1-GFP foci are absent in pre-mitotic nuclei marked by a hemispindle but predominantly expressed in nuclei that developed a short mitotic spindle, with PfARK1-GFP localizing at the spindle poles (Fig. 1B). Consistent with this observation, co-staining with antibodies against the centromeric histone variant CENH3 revealed peripheral association of PfARK1-GFP with the centromeric cluster specifically in nuclei displaying duplicated CPs (Fig. 1C).

**Figure 1.**
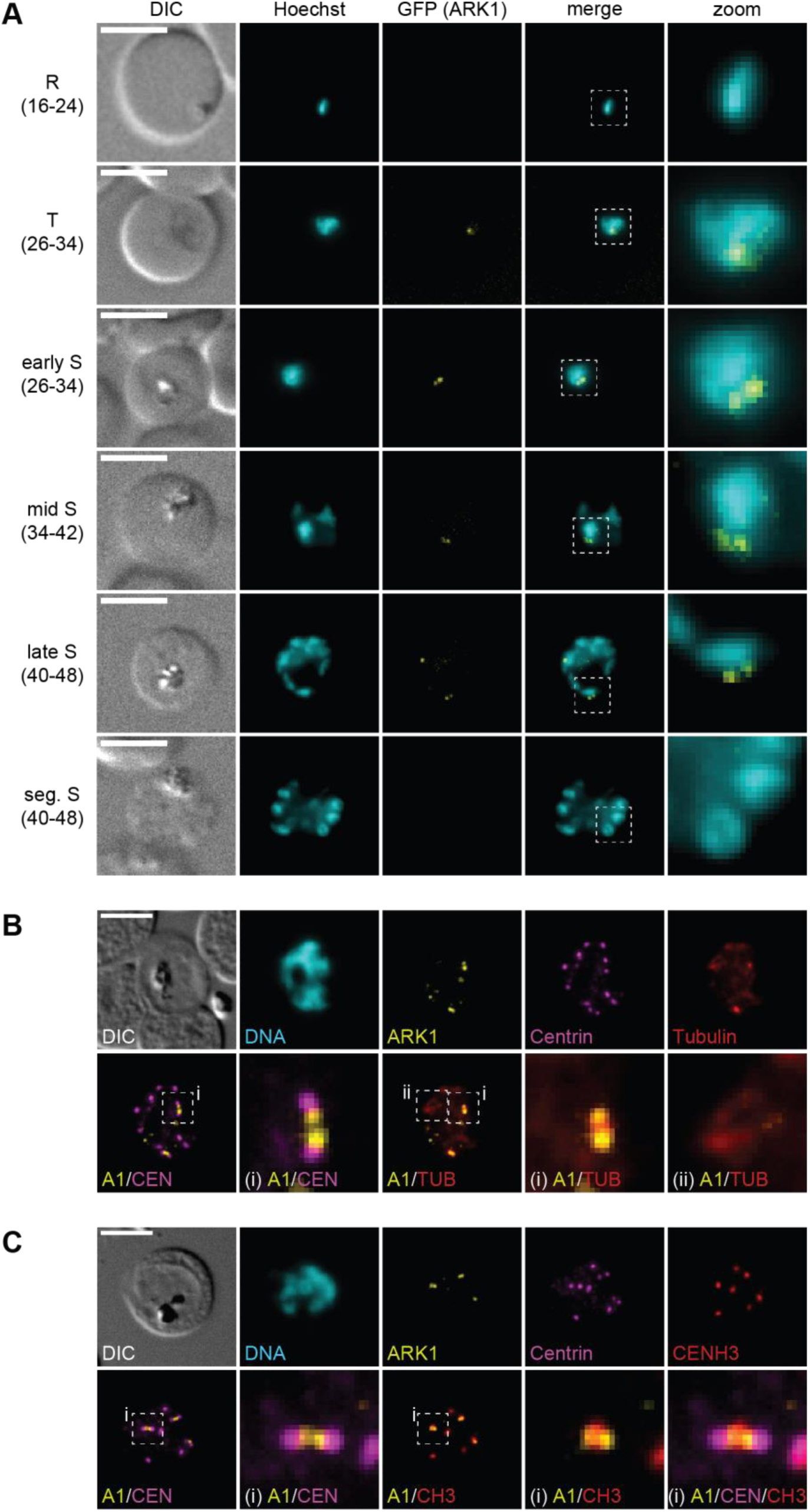
PfARK1 is expressed in a subset of nuclei during schizogony. **(A)** Localization of PfARK1-GFP throughout intra-erythrocytic asexual development as assessed by live cell fluorescence microscopy. Numbers in brackets indicate the age range (hours post invasion) of the synchronous parasite samples imaged at consecutive time points across intra-erythrocytic development. R, ring stage; T, trophozoite; S, schizont; seg., segmented. **(B-C)** Localization of PfARK1-GFP, Centrin and α/β- tubulin (B) or CENH3 (C) in schizonts as assessed by IFAs. Representative images are shown in each panel. DNA was stained with Hoechst (live) or DAPI (IFA). DIC, differential interference contrast. A1, PfARK1; CEN, Centrin; TUB, α/β-tubulin; CH3, CENH3; i/ii, zoom selections. Scale bar = 5 µm.

Similar to PfARK1, PfARK2-GFP localized to one or two distinct perinuclear foci first apparent in late trophozoites/early schizonts and then throughout schizogony until PfARK2-GFP signals disappeared in segmented schizonts, as assessed by live cell fluorescence microscopy (Fig. 2A). Furthermore, IFAs revealed that PfARK2-GFP also localizes directly adjacent to the outer CP marked by Centrin (Figs. 2B-C). In contrast to PfARK1, however, this association was observed in virtually all nuclei of developing schizonts (Figs. 2B-C). Triple-labeling IFAs co-staining with antibodies against Centrin and α/β-tubulin suggested that in early mitotic nuclei, PfARK2-GFP localizes to the spindle poles but not along the entire mitotic spindle (Fig. 2B). IFAs using α-CENH3 antibodies revealed a close association of PfARK2-GFP with the centromeric cluster in all nuclei throughout schizogony, indicating that PfARK2 may also localize to kinetochores (Fig. 2C).

**Figure 2.**
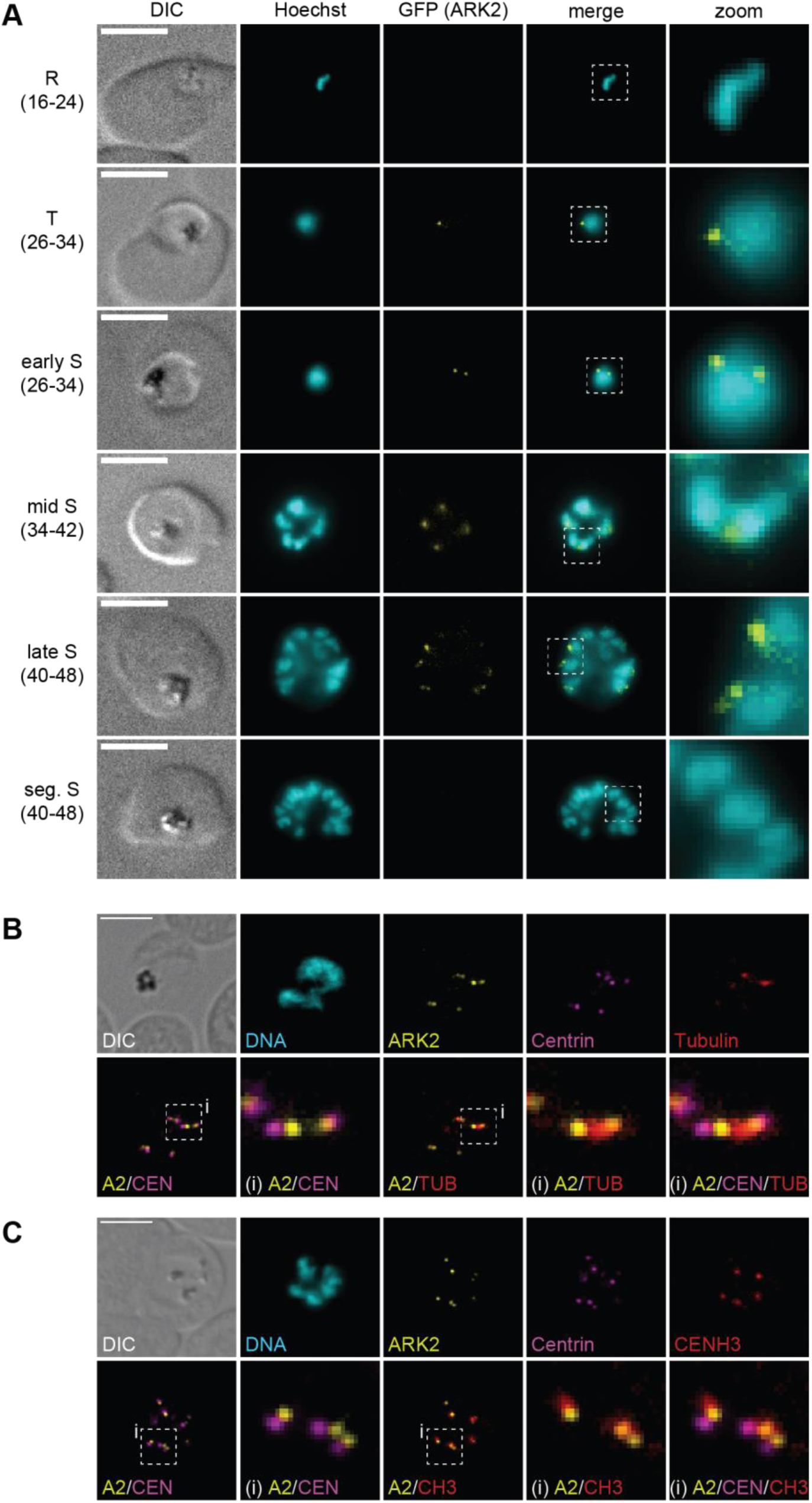
PfARK2 is expressed in all nuclei during schizogony. **(A)** Localization of PfARK2-GFP throughout intra-erythrocytic asexual development as assessed by live cell fluorescence microscopy. Numbers in brackets indicate the age range (hours post invasion) of the synchronous parasite samples imaged at consecutive time points across intra-erythrocytic development. R, ring stage; T, trophozoite; S, schizont; seg., segmented. **(B-C)** Localization of PfARK2-GFP, Centrin and α/β-tubulin (B) or CENH3 (C) in schizonts as assessed by IFAs. Representative images are shown in each panel. DNA was stained with Hoechst (live) or DAPI (IFA). DIC, differential interference contrast. A2, PfARK2; CEN, Centrin; TUB, α/β-tubulin; CH3, CENH3; i, zoom selections. Scale bar = 5 µm.

In line with previously published data (40), we detected PfARK3-GFP expression exclusively in late stage schizonts, marking distinct structures in close proximity to the Hoechst-stained area of individual nuclei (Fig. 3A). In segmented schizonts, PfARK3-GFP was again undetectable. The PfARK3-GFP signals observed in late stage schizonts appeared either as distinct single or paired foci or as elongated structures (Fig. 3A). Interestingly, IFAs suggested that these two types of localization patterns are mutually exclusive in individual schizonts, with all PfARK3-GFP signals being either of the focal or elongated type (Fig. 3B). Co-staining for Centrin revealed partial co-localization of the focal PfARK3-GFP signals with the outer CP from where the elongated PfARK3-GFP structures seem to emanate (Fig. 3B). Importantly, co-staining for CENH3 clearly demonstrated that in contrast to PfARK1-GFP and PfARK2-GFP, PfARK3-GFP localizes to the cytoplasmic side of the CP, which is further evident from the positioning of the elongated PfARK3-GFP structures in between adjacent nuclei in segmenting schizonts (Figs. 3B).

**Figure 3.**
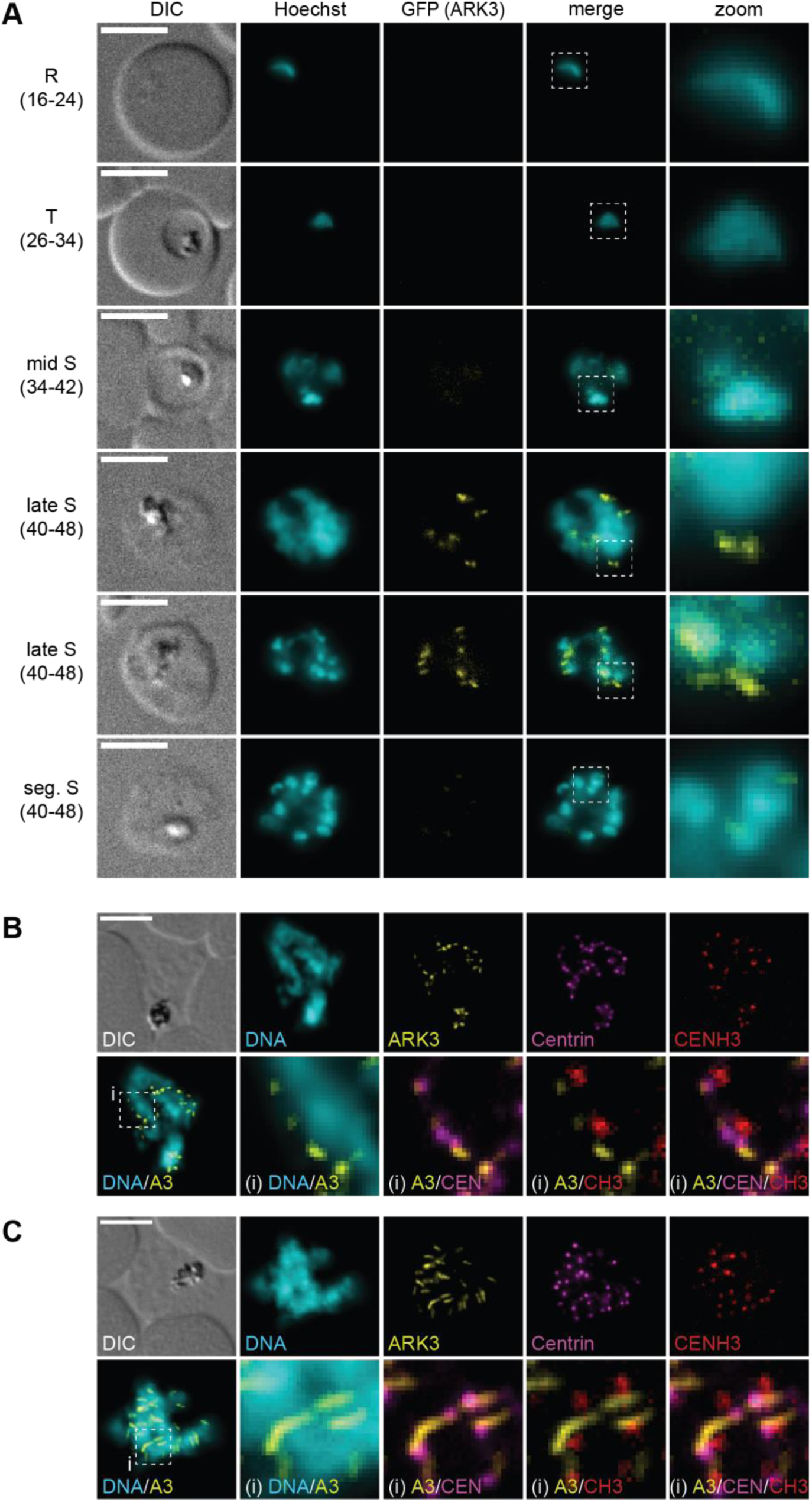
PfARK3 expression is restricted to late stage schizonts. **(A)** Localization of PfARK3-GFP throughout intra-erythrocytic asexual development as assessed by live cell fluorescence microscopy. Numbers in brackets indicate the age range (hours post invasion) of the synchronous parasite samples imaged at consecutive time points across intra-erythrocytic development. R, ring stage; T, trophozoite; S, schizont; seg., segmented. **(B-C)** Localization of PfARK3-GFP, Centrin and CENH3 in late stage schizonts as assessed by IFAs. Representative images are shown in each panel. DNA was stained with Hoechst (live) or DAPI (IFA). DIC, differential interference contrast. A3, PfARK3; CEN, Centrin; CH3, CENH3; i, zoom selections. Scale bar = 5 µm.

In summary, we show that the three Aurora-related kinases PfARK1-3 have unique expression dynamics during schizogony and display distinct spatial associations with the intranuclear body, mitotic spindle and centromeric cluster (PfARK1, PfARK2) or with the outer CP specifically in late stage schizonts (PfARK3).

### The DD/Shield system is unsuitable to investigate PfARK function

To study the function of the three PfARKs, we treated young ring stage parasites with RAPA to activate DiCre-mediated recombination, leading to replacement of the C-terminal GFP tag with the 3xHA-DD conditional knockdown tag (Fig. S1). DiCre-mediated recombination was confirmed in all three cell lines using PCR on gDNA (Fig. S1). While Shield-1 removal resulted in a moderate decrease of PfARK2-3xHA-DD expression levels in RAPA-treated parasites, PfARK1-3xHA-DD and PfARK3-3xHA-DD expression was not markedly reduced compared to parasites maintained in the presence of Shield-1 as shown by IFAs and Western blotting (Fig. S2).

Despite the ineffective knockdown of PfARK-3xHA-DD expression in RAPA-treated parasites in the absence of Shield-1, we still investigated whether potential knockdown phenotypes could be observed. However, parasite multiplication rates of RAPA-treated parasites cultured in the absence of Shield-1 did not differ from DMSO-treated control parasites (expressing PfARK-GFP) over two subsequent invasion cycles as assessed by flow cytometry (Fig. S2). Based on the observed co-induction of *pfark2* and *pfap2-g* transcription in lysoPC-depleted parasites (8), we were also interested to examine whether the moderate knockdown of PfARK2-3xHA-DD expression achieved in the absence of Shield-1 may have any effect on parasite SCRs. To this end, we compared the SCRs between RAPA-treated PfARK2-3xHA-DD knockdown parasites cultured in the absence of Shield-1 and DMSO-treated control parasites expressing PfARK2-GFP, cultured for one cycle in minimal fatty acid medium (mFA) supplemented with 2 mM choline chloride (commitment-suppressing conditions) and mFA medium lacking choline chloride (commitment-inducing conditions) (8). SCRs were assessed by quantifying the proportion of sexually committed PfAP2-G-mScarlet-positive ring stage progeny among all ring stage parasites using high content imaging as previously described (55). As shown in Fig. S2, we observed no difference in SCRs between parasites cultured under protein knockdown and control conditions, neither for the NF54/ARK2-GFP-icKD line nor for the NF54/ARK1-GFP-icKD and NF54/ARK3-GFP-icKD lines that were also tested.

In summary, we were unable to observe any defects on asexual parasite development, multiplication or sexual commitment using the DD-based approach as an attempt to conditionally knock down PfARK expression. Alternative conditional knockdown approaches such as the *glmS* riboswitch (56) or TetR-DOZI-aptamer systems (57) or a DiCre-based *pfark* knockout strategy will therefore be required to study PfARK function in *P. falciparum*.

### PfARK expression and localization in gametocytes

Next, we monitored PfARK expression and localization throughout gametocyte development by live cell fluorescence microscopy. While PfARK1-GFP and PfARK2-GFP expression was readily detected (Figs. 4 and 5), PfARK3-GFP signals were not observed in gametocytes (Fig. S3). PfARK1-GFP was not expressed at appreciable levels in stage I to III gametocytes (day 2-6 of gametocyte development), but was detectable in stage IV and V gametocytes (days 8-12 of gametocyte development), showing weak PfARK1-GFP signals throughout the Hoechst-stained area of the nucleus and a distinct and more densely stained spot (Fig. 4A). Interestingly, PfARK1-GFP expression was only detected in a subset of late stage gametocytes, suggesting this kinase may be expressed in a sex-specific manner. We therefore performed dual-labeling IFAs co-staining for PfARK1-GFP and Pfg377, a female-enriched osmiophilic body protein routinely used as a marker to distinguish between female and male late stage gametocytes (58–62). Indeed, automated quantification of fluorescence signal intensities revealed mutually exclusive expression for PfARK1-GFP and Pfg377 in stage V gametocytes, indicating that PfARK1 is specifically expressed in male gametocytes (Fig. 4B-C).

**Figure 4.**
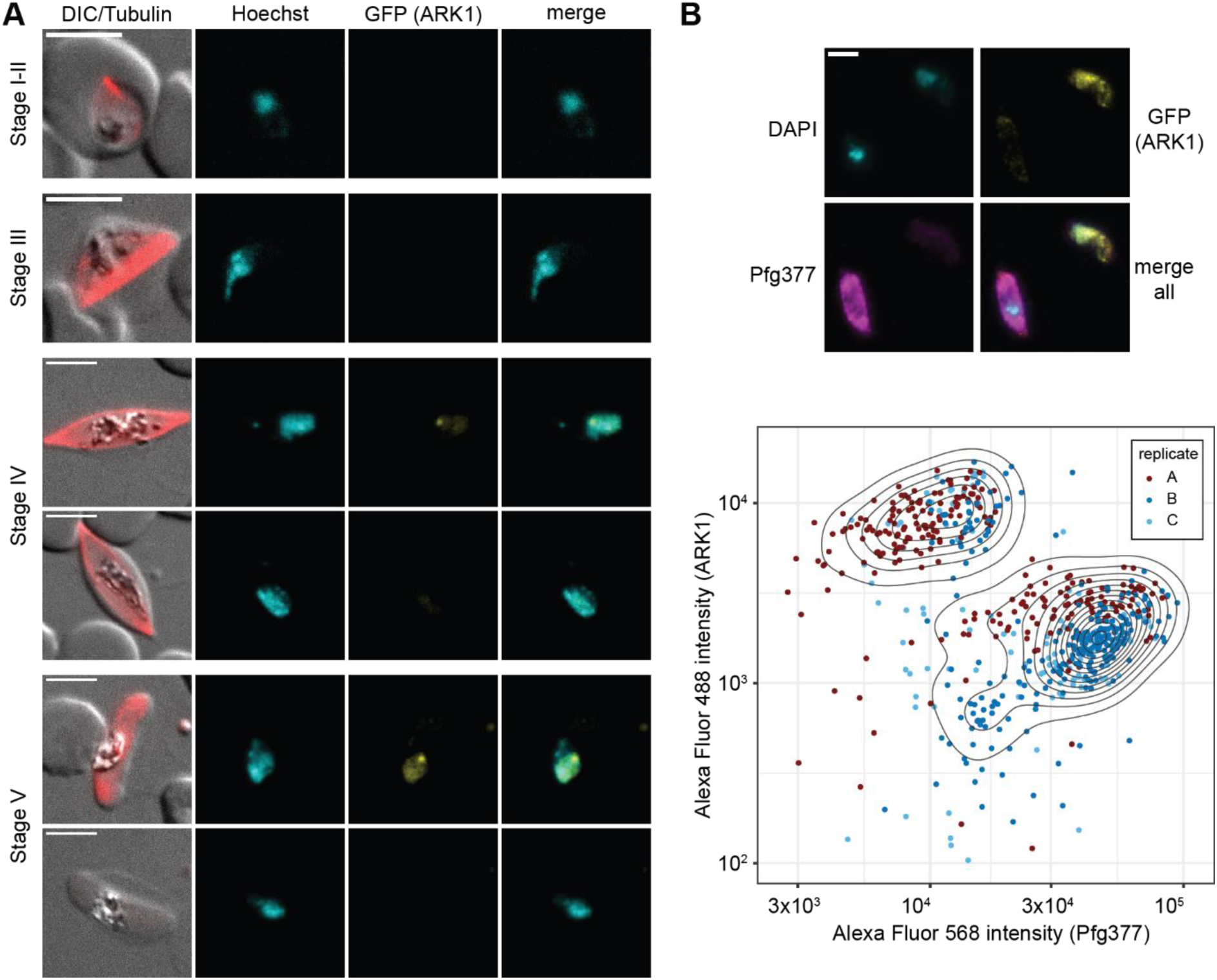
PfARK1 shows male-specific expression in late stage gametocytes. PfARK1-GFP expression in gametocytes was assessed by live cell fluorescence microscopy. Representative images are shown. DNA was stained with Hoechst. MTs were stained with SPY555-tubulin. Stages of gametocyte development are indicated. Representative images are shown. DIC, differential interference contrast. Scale bar = 5 µm. **(B)** Top: Localization of PfARK1-GFP (yellow) and Pfg377 (magenta) in stage V gametocytes as assessed by IFA. Representative images are shown. DNA was stained with DAPI. Scale bar = 5 µm. Bottom: Quantification of PfARK1-GFP (Alexa Fluor 488) and Pfg377 (Alexa Fluor 568) IFA fluorescence signal intensities in single stage V gametocytes. The assay was performed in three independent biological replicates (251, 259 and 137 cells analyzed per replicate). Dots in the contour plot represent the fluorophore intensities measured for individual cells.

**Figure 5.**
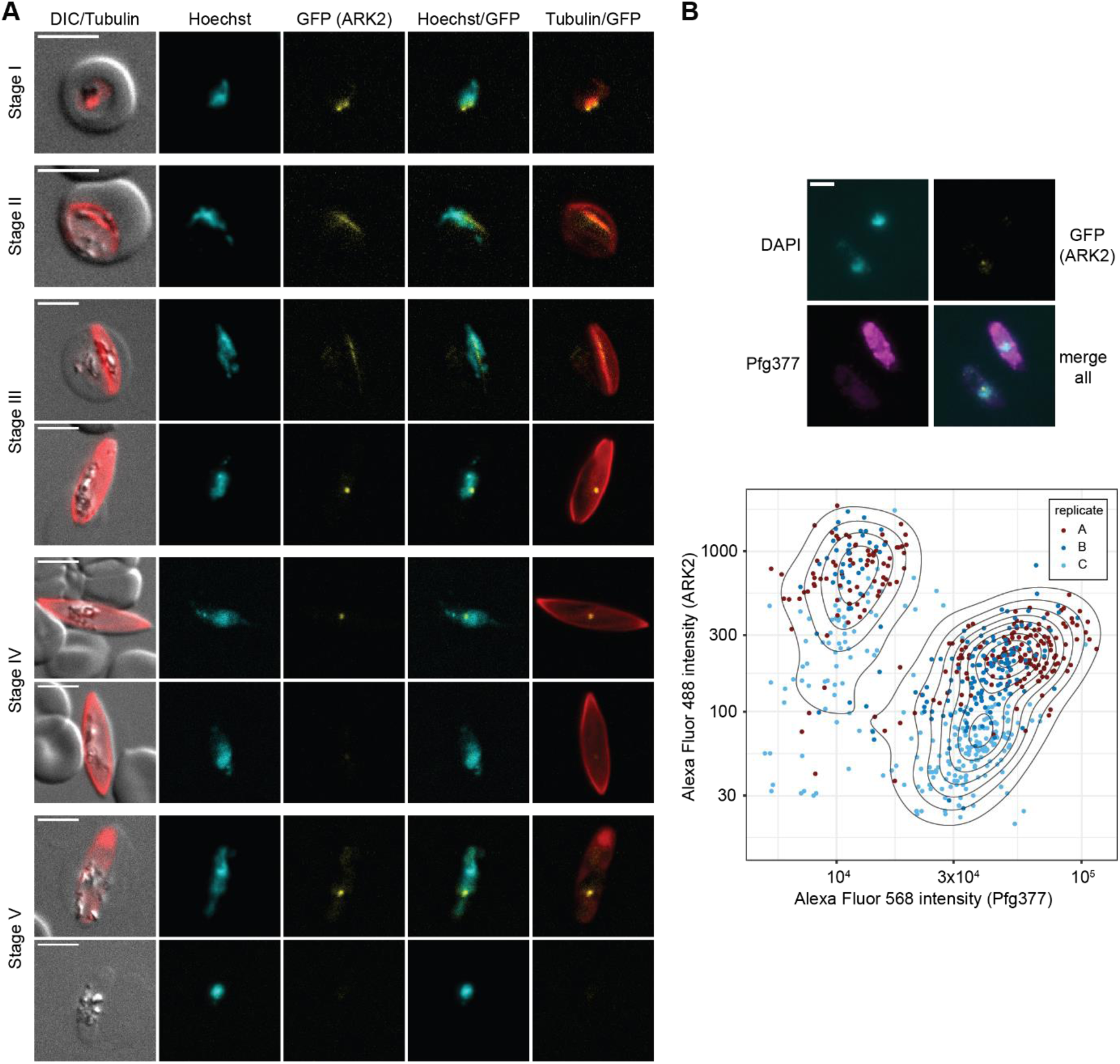
PfARK2 is expressed throughout gametocytogenesis with male-specific expression observed in late stage gametocytes. **(A)** PfARK2-GFP expression in gametocytes was assessed by live cell fluorescence microscopy. Representative images are shown. DNA was stained with Hoechst. MTs were stained with SPY555-tubulin. Stages of gametocyte development are indicated. Representative images are shown. DIC, differential interference contrast. Scale bar = 5 µm. **(B)** Top: Localization of PfARK2-GFP (yellow) and Pfg377 (magenta) in stage V gametocytes as assessed by IFA. Representative images are shown. DNA was stained with DAPI. Scale bar = 5 µm. Bottom: Quantification of PfARK2-GFP (Alexa Fluor 488) and Pfg377 (Alexa Fluor 568) IFA fluorescence signal intensities in single stage V gametocytes. The assay was performed in three independent biological replicates (231, 170 and 242 cells analyzed per replicate). Dots in the contour plot represent the fluorophore intensities measured for individual cells.

In contrast to PfARK1-GFP, PfARK2-GFP was expressed throughout sexual development with signals detectable in all five gametocyte stages (Fig. 5A). In stage I gametocytes (day 2), PfARK2-GFP localized to two distinct foci at the nuclear periphery, partially co-localizing with nuclear MTs stained with the live cell-compatible SPY555-tubulin stain. In stage II and III gametocytes, PfARK2-GFP specifically co-localized along the elongated bundle of nuclear MTs that span the entire nucleus (22) (Fig. 5). Coincident with the disappearance of the intranuclear microtubular structure in stage IV gametocytes, PfARK2-GFP accumulated at a single densely stained focus that remained visible until the end of gametocytogenesis. Similar to PfARK1, we again observed a PfARK2-GFP-negative population from stage IV gametocytes onwards, suggesting sex-specific expression in late stage gametocytes. Dual-labeling IFAs co-staining for PfARK2-GFP and Pfg377 confirmed that PfARK2-GFP expression in late stage gametocytes is also male-specific. Together, these findings show that during the early phase of gametocyte differentiation PfARK2 is expressed in all gametocytes, whereas from stage IV onwards only male gametocytes continue to express PfARK2 (Fig. 5 B-C).

Based on our observations made in asexual parasites, we reasoned that the punctate region marked by PfARK1-GFP and PfARK2-GFP in late stage male gametocytes may correspond to the inner CP. IFAs co-staining for PfARK1-GFP/PfARK2-GFP and Centrin in stage IV/V gametocytes demonstrated that indeed both kinases localize at the nuclear periphery adjacent to the outer CP (Fig. 6A-B). These experiments further confirmed that PfARK1-GFP and PfARK2-GFP were only expressed in a subset of cells, in contrast to Centrin that was observed in all gametocytes (Fig. 6A-B). Interestingly, in male gametocytes we routinely observed two closely paired Centrin foci directly adjacent to the perinuclear PfARK1-GFP/PfARK2-GFP signals. In contrast, female gametocytes, identified based on the lack of PfARK1-GFP/PfARK2-GFP expression, consistently displayed only a single Centrin focus (Fig. 6A-B).

**Figure 6.**
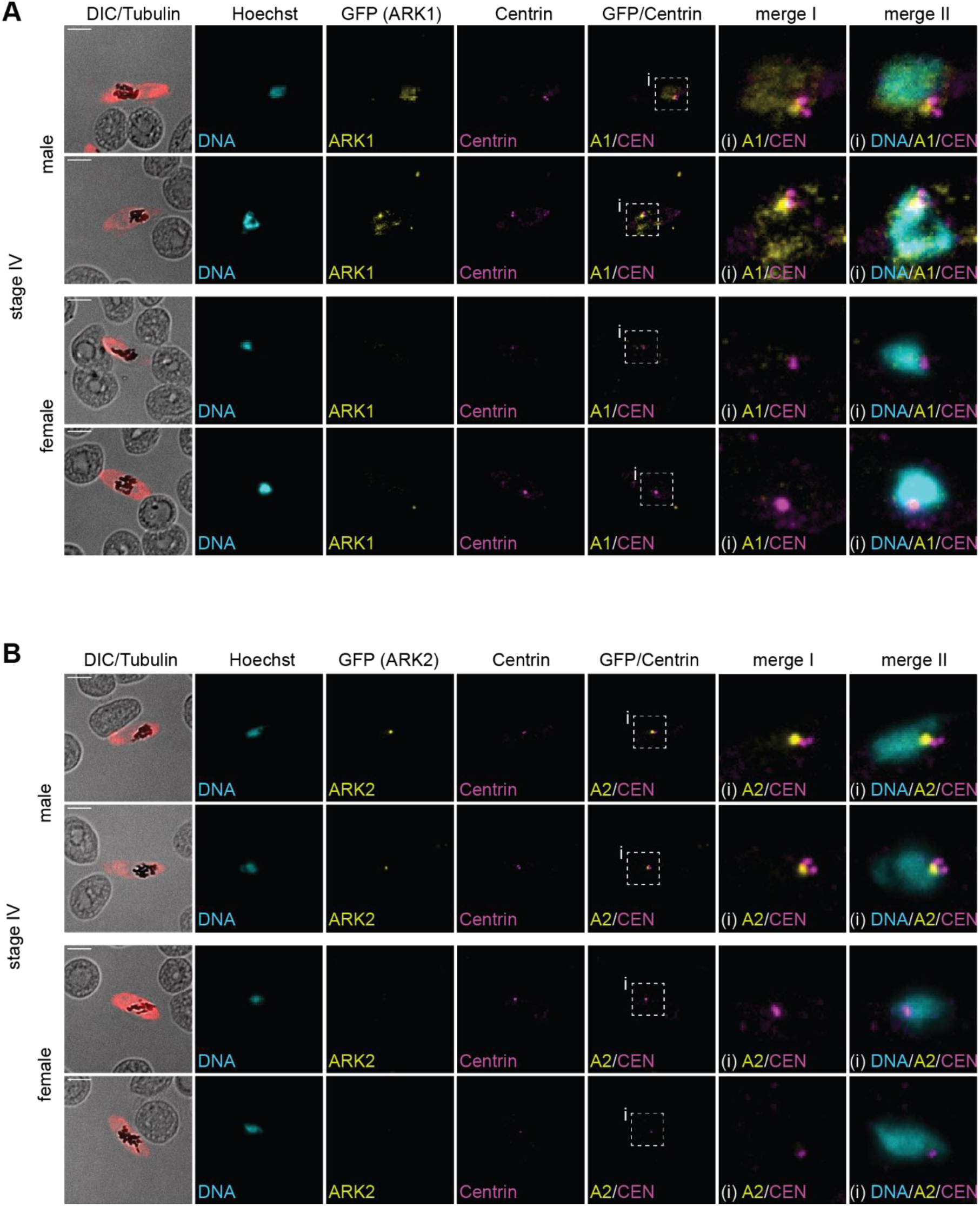
PfARK1 and PfARK2 localise to the intranuclear body in male late stage gametocytes. **(A-B)** Localization of PfARK1-GFP and Centrin (A) or PfARK2-GFP and Centrin (B) in female and male stage IV gametocytes as assessed by IFAs. MTs were stained with antibodies against α/β- tubulin. Representative images are shown in each panel. DNA was stained with DAPI. DIC, differential interference contrast. A1/2, PfARK1/PfARK2; CEN, Centrin; i, zoom selections. Scale bar = 5 µm.

In summary, this set of experiments demonstrated distinct expression timing, subcellular localization and sex-specific expression for PfARK1 and PfARK2 during gametocytogenesis, while PfARK3 expression was not detected. PfARK1 is only expressed in late stage male gametocytes and localizes to a perinuclear spot directly adjacent to the outer CP, and more diffusely to the chromatin-rich region of the nucleus. PfARK2 initially localizes along the nuclear MTs in all stage I-III gametocytes before adopting male-specific expression in stage IV and V gametocytes, where PfARK2 also shows exclusive localization to a single focus at the nuclear periphery in direct vicinity to the outer CP, similar to PfARK1.

## Discussion

In this study, we set out to further characterize the expression, subcellular localization and function of the three Aurora-related kinases PfARK1, PfARK2 and PfARK3 in *P. falciparum* asexual and sexual blood stage parasites. Unfortunately, our attempts to study PfARK function using an inducible conditional knockdown approach relying on the DD/Shield-1 system failed due to inefficient reduction of PfARK-3xHA-DD protein levels. While the exact underlying cause remains unclear, the degradation efficiency of DD-tagged proteins is somewhat unpredictable and dependent on proteasome accessibility (53, 63). Previous studies further indicated that for proteins with enzymatic activities, including several parasite kinases, highly efficient knockdown of expression may be required to observe loss-of-function phenotypes (64–67), offering a possible explanation for why the moderate reduction of PfARK2-3xHA-DD levels in absence of Shield-1 had no effect on parasite multiplication. In this context, it is noteworthy that conditional depletion of PbARK2 in *P. berghei* using the auxin-inducible degron system was also unsuccessful (36). Clearly, conditional expression systems allowing for a more efficient knockdown of protein/mRNA expression levels, or even DiCre-dependent inducible gene knockouts, will be required to study PfARK function.

We found that all three PfARKs are expressed in asexual parasites exclusively during schizogony but with distinct temporal expression and localization patterns. Our results on PfARK1 are in line with and expand on the data previously published by Reininger and colleagues (38, 39). PfARK1 localized to a single or two closely adjacent perinuclear foci first observed in single-nucleated parasites that likely represented early schizonts in their first S-phase or at the onset of mitosis. Throughout schizogony, PfARK1 was only expressed in a subset of nuclei, the majority of which were early mitotic nuclei marked by a duplicated CP and short mitotic spindle and the two PfARK1 foci localized internally to both outer CPs flanking the intervening spindle or centromeric cluster. In contrast, PfARK1 signals were absent in pre-mitotic nuclei characterised by a hemispindle and in most DAPI-stained areas marked by a single or well separated CPs that we believe corresponded to nuclei at early S-phase or anaphase/late mitosis, respectively. Based on this periodic expression behaviour and distinct localization pattern, we conclude that PfARK1 transiently localizes to the intranuclear body and spindle poles specifically from late S-phase until the metaphase of mitosis. However, we cannot exclude the possibility that PfARK1 also (or instead) localizes to the centromeres/kinetochores because our wide-field microscopy data lack the resolution required to visualise the closely adjacent intranuclear body and centromeric cluster as separate entities.

During gametocytogenesis, PfARK1 showed no discernible expression in stage I to III gametocytes. Interestingly, during this phase of gametocyte development the centromeres do not form a single cluster but are laterally sequestered along the bundle of nuclear MTs, as evidenced by co-staining for the outer kinetochore marker NDC80 or CENH3 (22). The lack of PfARK1 expression in these cells therefore provides evidence that PfARK1 is not required for kinetochore-MT attachment. In stage IV/V gametocytes, where the nuclear MTs have been disassembled and centromeres again conjoin into a single cluster in close proximity to the CP (22), we found that PfARK1 was expressed and localized directly adjacent to the outer CP (presumably to the intranuclear body and/or centromeric cluster) exclusively in male gametocytes. This sex-specific expression is consistent with an important function of PfARK1 as a regulator of mitosis since only male gametocytes, but not females, will undergo mitotic cell division upon activation in the mosquito midgut. Interestingly, we consistently observed two closely paired Centrin foci specifically in the PfARK1-positive subpopulation, indicative for separation of the outer CP in male, but not in female late stage gametocytes. This finding is in line with those recently published by Li and colleagues who observed two adjacent Centrin-4 foci in about 15% of late stage gametocytes and speculated these cells may represent male gametocytes (22). The expression of PfARK1 and duplication of the outer CP already at stage IV of sexual development, i.e. several days before male gametocytes developed into transmissible stage V gametocytes capable of entering S-phase and mitosis, is noteworthy and suggests that male gametocytes prepare early to coordinate the three rapid rounds of endomitosis and axoneme assembly during male gametogenesis.

Collectively, the spatiotemporal expression pattern of PfARK1 during schizogony and gametocyte differentiation suggests that PfARK1 may primarily regulate CP duplication and/or maturation and nucleation of the bipolar mitotic spindle, akin to the functions known for mammalian Aurora A. Furthermore, the apparent absence of PfARK1 in the later phase of mitosis is reminiscent of Aurora A degradation during anaphase and mitotic exit. Aurora A degradation is mediated by the anaphase promoting complex/cyclosome (APC/C), a multi-subunit ubiquitin ligase that targets key regulators of mitosis for proteasomal degradation to control cell cycle progression (37, 68). Although malaria parasites encode only a largely reduced set of recognizable APC/C components (68, 69), all of which seem to be essential for parasite proliferation (69), it is tempting to speculate that PfARK1 degradation may similarly be mediated by APC/C-dependent ubiquitination. Despite these apparent analogies between PfARK1 and the distantly related mammalian Aurora A, the direct PfARK1 ortholog of *T. gondii*, TgARK1, shows a markedly different localization pattern and has been proposed to represent a functional homolog of Aurora B (70). *T. gondii* tachyzoites divide by endodyogeny, which typically entails a single round of S-phase and mitosis followed by daughter cell budding inside the mother cell (11). While to our knowledge TgARK1 localization has not been assessed in conjunction with known markers of mitotic structures, Berry and colleagues reported that TgARK1 localizes throughout the nucleus during G1-phase and to the nucleus and cytoplasm during cell division (70). Furthermore, TgARK1 co-purified with TgINCENP1 and mislocalized upon depletion of TgINCENP2, two putative homologs of the human CPC component INCENP that is known to interact with Aurora B. Importantly, conditional overexpression of a TgARK1 dead kinase mutant resulted in kinetochore detachment from centromeres and prevented faithful chromosome segregation, consistent with Aurora B-like functions at the kinetochore-centromere interface (70). However, these dominant negative mutants also exhibited severe defects in centrosome inner core separation, maturation of the spindle pole bodies (centrocones) and nuclear mitotic spindle, suggesting that TgARK1 may also possess Aurora A-like activities in regulating the early steps of mitosis (70).

The subcellular localization of PfARK2 in asexual parasites was similar to that observed for PfARK1. PfARK2 localized to one or two perinuclear foci in single-nucleated late trophozoites/early schizonts and throughout schizogony, consistent with recent observations made for PbARK2 in *P. berghei* (36). In early mitotic nuclei, PfARK2 localized directly internal to both outer CPs and at the periphery of the mitotic spindle/centromeric cluster. In striking difference to PfARK1, however, PfARK2 was expressed in all nuclei of developing schizonts, irrespective of mitotic stage, and consistently localized in between the outer CP and the closely associated centromeric cluster. While we are unable to discriminate between a localization to the intranuclear body or kinetochore-loaded centromeres, we assume that PfARK2 localizes to both structures. First, PfARK2 localized along the intranuclear MTs in stage I-III gametocytes, which provides at least circumstantial evidence that PfARK2 may be involved in kinetochore-MT attachment during this phase of sexual differentiation. Second, in stage IV/V gametocytes, PfARK2 displayed the same sex-specific expression and specific localization in close juxtaposition to a duplicated outer CP as observed for PfARK1. Third, during male gametogenesis in *P. berghei*, PbARK2 is closely associated with the inner CP and laterally distributed along the mitotic spindles internal to and partially overlapping with NDC80, suggesting localization at the spindle-kinetochore interface (36). Importantly, reciprocal immunoprecipitation of crosslinked PbARK2 and PbEB1 complexes followed by protein mass spectrometry confirmed the close association of PbARK2 with various kinetochore components including NDC80, NUF2 and the putative CPC protein INCENP2, MT-associated proteins such as PbEB1, myosin K and kinesin 8X as well as cohesin and condensin subunits (36). Besides the well-described interaction with INCENP, kinesins, EB1 and cohesion-/condensin-associated factors are also known interactors/substrates of Aurora B (37). Together, these results indicate that PfARK2/PbARK2 are primarily involved in spindle dynamics and in regulating kinetochore/centromere attachment to the nuclear spindle and chromosome segregation, thereby sharing at least some of the functions known for Aurora B. As a further notion, TgARK2 was also reported to localize to the spindle pole body in late G1-phase and to associate with the mitotic spindle and kinetochores during mitosis in tachyzoites (40), suggesting that PfARK2/PbARK2 and TgARK2 are functional orthologs. However, while PfARK2 and PbARK2 are considered essential for mitotic proliferation of malaria blood stage parasites, TgARK2 is dispensable for tachyzoite replication and growth (40). The reason(s) underlying this discrepant requirement for ARK2 function are unknown but may be related to the different modes of mitotic cell division employed by these parasite stages (schizogony vs endodyogeny) or functional redundancy between TgARK2 and another kinase, for instance TgARK1.

Similar to previously published results (40), we found that PfARK3 was only expressed in multi-nucleated late stage schizonts, where it was confined to small foci or rod-like structures in close proximity to the nuclei. Interestingly, these two distinct PfARK3 localization patterns were essentially mutually exclusive, such that in individual schizonts all PfARK3 signals were either of a punctate or markedly elongated type. Co-staining for Centrin and CENH3 revealed that in schizonts exhibiting the punctate signal, PfARK3 partially overlapped with the cytoplasmic side of the outer CPs. In schizonts displaying the elongated type, PfARK3 stretched away from the outer CPs in shapes reminiscent of the SPMTs that develop in nascent merozoites during the segmentation process (71, 72) and also seem to emanate from the outer CP (18, 22). In segmented schizonts, PfARK3 signals were again undetectable. These results indicate that PfARK3 does not participate in mitosis but rather in the coordination of synchronous cell division at the end of schizogony. The lack of PfARK3 expression in the non-replicative gametocyte stages provides additional support for this hypothesis. Moreover, our data on PfARK3 are consistent with the results obtained for TgARK3 in dividing *T. gondii* tachyzoites (40, 73). In these stages, TgARK3 also localizes to the outer core of the duplicated centrosomes during S-phase and then elongates along one side of the emerging cytoskeleton of the budding daughter parasites as they progress through mitosis and cytokinesis (40, 73). Interestingly, treatment with the microtubule-depolymerising small molecule oryzalin showed that TgARK3 localization is indeed dependent on intact SPMTs (73). Furthermore, TgARK3 knockout parasites are inviable, and phenotyping of a conditional knockdown mutant revealed that TgARK3 function is not required for completion of mitosis, but for proper daughter cell budding, cytokinesis and parasite growth (40). Together, these data indicate that PfARK3 and TgARK3 are functional orthologs and highly divergent members of the Aurora kinase family that underwent functional adaptation to control cell division during schizogony and endodyogeny, respectively. While their exact role(s) in these processes remain to be determined, the specific spatiotemporal expression patterns of PfARK3 and TgARK3 point at crucial functions in daughter cell cytoskeleton assembly and dynamics, as suggested earlier (40, 73).

In conclusion, our study presents new insight into the expression dynamics and subcellular localization of all three *P. falciparum* ARKs throughout schizogony and gametocyte development. We describe distinct temporal and spatial associations for each PfARK with mitotic structures that are clearly indicative for essential non-overlapping roles during mitosis (PfARK1, PfARK2) and cytokinesis (PfARK3) and offer testable hypotheses for their targeted functional analysis. Furthermore, the transgenic lines generated here provide excellent tools for future research – including super-resolution microscopy techniques to determine PfARK localization at high resolution, protein interactome studies to identify PfARK binding partners/substrates and further genetic modifications to study PfARK function – to understand in molecular detail how these divergent mitotic kinases regulate cell cycle progression in malaria parasites.

## Methods

### P. falciparum cell culture

*P. falciparum* NF54 parasites were cultured as described previously (74) in AB+ or B+ human red blood cells (Blood Donation Center, Zürich, Switzerland) at a hematocrit of 5% in medium containing 10.44 g/l RPMI-1640, 25 mM HEPES, 100 μM hypoxanthine, 24 mM sodium bicarbonate, 0.5% AlbuMAX II (Gibco #11021-037) and 0.1 g/l neomycin. For routine parasite propagation, 2 mM choline chloride (Sigma #C7527) was added to the culture medium to suppress sexual commitment (8). Parasites were synchronized using consecutive sorbitol treatments (75). To induce the DiCre-mediated recombination of loxP sites, early ring stage parasites (0-6 hpi) were treated for 4 hours with 100 nM rapamycin (49). To achieve stabilization or destabilization of DD-tagged proteins, parasites were cultured in the presence or absence of 625 nM Shield-1, respectively (52, 53). Cultures were gassed with 3% O_2_, 4% CO_2_ and 93% N_2_ and incubated at 37 °C.

To obtain synchronous gametocyte cultures, sexual commitment was induced in by exposing synchronized ring stage cultures (16-24 hpi, 1-1.5% parasitemia) to minimal fatty acid medium (mFA) for 32 hours as described (8). The mFA medium contains 0.39% fatty acid-free BSA (Sigma #A6003) instead of 0.5% AlbuMAX II and 30 μM each of oleic and palmitic acid (Sigma #O1008 and #P0500). After completion of the invasion cycle, the ring stage progeny was cultured for six days in medium containing 10% human serum (Blood Donation Centre, Basel, Switzerland) and supplemented with 50 mM N-acetyl-D-glucosamine (GlcNAc) to eliminate asexual parasites (76). From day seven onwards, gametocytes were cultured in absence of GlcNAc and daily medium changes were performed on a 37 °C heating plate.

### Cloning of transfection constructs

Gene editing of NF54 parasites was performed using a CRISPR/Cas9 approach based on the co-transfection of two plasmids as previously described (9). The first pHF-gC-derived plasmid contains expression cassettes for the *S. pyogenes* Cas9 endonuclease, the single guide RNA (sgRNA) and the resistance marker human dihydrofolate reductase. The second pD-derived donor plasmid contains the template for homology-directed repair of the Cas9-induced DNA double-strand break (Fig. S1). The gene editing strategies for *pfark1* (PF3D7_0605300), *pfark2* (PF3D7_0309200) and *pfark3* (PF3D7_1356800) were identical. Therefore, the following description of plasmid generation applies to all three target genes, with “*x*” referring to the *pfark* variant. To obtain plasmid pHF-gC_ARKx, the sequence encoding a sgRNA targeting the 3’ end of *arkx* was assembled by annealing two complementary oligonucleotides (gRNA_ARKx_a and gRNA_ARKx_b) and cloned into the BsaI-digested pHF-gC backbone (9) using T4 DNA ligase (New England Biolabs #M0202). The donor plasmid pD_ARKx_icKD was produced by assembling four DNA fragments in a Gibson Assembly reaction (77) using (i) the plasmid backbone amplified by PCR from pUC19 (primers PCR1_F/PCR1_R) (9), (ii/iii) the 5’ and 3’ homology regions amplified from genomic DNA (gDNA) (primers PCR2_ARKx_F/PCR2_ARKx_R and PCR3_ARKx _F/PCR3_ARKx _R, respectively), and (iv) a sequence assembly containing a loxP intron element (loxPint) (50), the *gfp* coding sequence, the *hrpII 3’* terminator sequence, a second loxPint element and a sequence encoding three hemagglutinin epitopes (3xHA) fused to the FKBP/DD destabilization domain (DD) (Fig. S1), amplified from a synthetic sequence ordered from Genscript (primers PCR4_ARKx _F/PCR4_ARKx _R). Primer sequences are listed in Table S1.

### Transfection and selection of genetically modified parasites

Transfection and selection of genetically modified parasites were performed as previously described (9). In brief, 50 μg of each plasmid (pHF-gC_ARKx and pD_ARKx_icKD) were co-transfected into NF54/DiCre parasites (54), in which PfAP2-G has previously been tagged with the fluorophore mScarlet (55). To select for transgenic parasites, 5 nM WR99210 (WR) was added 24 hours after transfection for six days, followed by continuous culture in the absence of WR until a stably propagating population was obtained. Successful gene editing was confirmed by PCR on genomic DNA (gDNA) (Fig. S1). Primer sequences are listed in Table S1.

### Western blotting

Parasites at 3-5% parasitaemia were released from infected RBCs by saponin lysis of 500 µl packed RBCs for 10 min on ice in 3 ml ice-cold 0.15% saponin in PBS. Parasite pellets were washed twice in ice-cold PBS and resuspended in a urea/SDS lysis buffer (8 M urea, 5% SDS, 2 mM EDTA, 1 mM TCEP, 50 mM Bis-Tris, and 1x protease inhibitor cocktail (Roche #11697498001), pH 6.5). Proteins were separated on 3-8% Tris-Acetate gels (Invitrogen #EA0375) using Tris-Acetate SDS running buffer (Invitrogen #LA0041). Proteins were transferred to a nitrocellulose membrane (GE Healthcare #106000169) followed by blocking with 4% skim milk in PBS (phosphate-buffered saline)/0.1% Tween (PBS-T) for 1 hour. The membrane was probed using the primary antibodies mAb rat α-HA (1:2,000, Roche Diagnostics #11867423001) or mAb mouse α-PfGAPDH (1:20,000) (78) diluted in blocking buffer. After a 2-hour incubation, the membrane was washed five times in PBS-T before it was incubated with the secondary antibody goat α-rat IgG (H&L)-HRP (1:10,000, Southern Biotech #3050-05) or goat α-mouse IgG (H&L)-HRP (1:10,000, GE healthcare #NXA931), respectively. After a 1-hour incubation, the membrane was washed five times and the signal was detected using the chemiluminescent substrate SuperSignal West Pico Plus (Thermo Scientific #34580) in conjunction with a Vilber Fusion FX7 Edge 17.10 SN imaging system.

### Staining of cells for live cell fluorescence microscopy

RBC suspensions at 2-5% parasitaemia were stained using 5 µg/ml Hoechst (Merck # 94403) for 20 min at 37 °C protected from light to stain the DNA. Tubulin staining was performed with SPY555-tubulin (Lubio Science #SC203) for 1 hour at 37 °C as recommended by the manufacturer. Stained cells were washed once in PBS and mounted on microscopy slides using VectaShield (Vector Laboratories #H-1200).

### Immunofluorescence assays

Thin blood smears were prepared and RBCs were fixed in ice-cold methanol or methanol/acetone (60:40) for 2 min. The fixed cells were incubated for 1 hour with blocking solution (3% BSA in PBS) followed by 1 hour incubation with the following primary antibodies diluted in blocking solution: mouse IgG1 mAb α-GFP (1:500, Roche Diagnostics #11814460001), mouse IgG2 mAb α-Centrin clone 20H5 (1:500, Sigma #ZMS1054) (21, 79); rabbit α-Pfg377 (1:1,000) (58); rabbit polyclonal α- CENH3 (1:500) (80); α-α/β-tubulin (1:500/1:1000, Geneva Antibody Facility #AA345/#AA344) (30). After three washes with PBS, the following secondary antibodies were incubated for 1 hour in blocking solution and protected from light: Alexa Fluor 488-conjugated α-mouse IgG (1:500, Molecular Probes #A11001), Alexa Fluor 488-conjugated α-mouse IgG1 (1:500, Molecular Probes #A21121), Alexa Fluor 568-conjugated α-mouse IgG2a (1:500, Molecular Probes #A21134), Alexa Fluor 568-conjugated α-rabbit IgG (1:500, Molecular Probes #A11011). After five washes in PBS, slides were air-dried for 10 min, 2 µl of Vectashield containing DAPI stain (Vector Laboratories #H-1200-10) was added and a coverslip mounted onto the slide.

### Fluorescence microscopy

Parasites were imaged using a Leica DM5000 B fluorescence microscope (40x and 63x objectives) equipped with a Leica K5 sCMOS camera and the Leica application suite software (LAS X Version: 3.7.5.24914). Image processing was performed with Fiji (ImageJ version 1.52n). Within each experiment, identical settings were used for both image acquisition and processing.

### Quantification of parasite multiplication rates

To assess the effect of DD-mediated degradation of PfARK-3xHA-DD expression on parasite multiplication, parasite cultures synchronized to the young ring stage (0-6 hpi, 0.2% parasitaemia) were split and treated for 4 hours with 100 nM rapamycin or DMSO (solvent control), respectively. Parasites were cultured for five consecutive days with daily medium changes. Parasitaemia was measured by flow cytometry after the first and second invasion cycles (generation 2, day 3; generation 3, day 5). Samples were stained with 1x SYBR Green DNA stain (Invitrogen #S7563), incubated for 20 min in the dark, and washed in PBS. Using a MACS Quant Analyzer 10 (Miltenyi Biotec), 50,000 events per condition were measured and analyzed using the FlowJo_v10.6.1 software. The proportion of infected RBCs was determined based on the SYBR Green signal intensity. The gating strategy is shown in Fig. S4.

### Quantification of sexual commitment rates

To assess the effect of DD-mediated degradation of PfARK-3xHA-DD expression on sexual commitment rates (SCRs), parasite cultures synchronized to the young ring stage (0-6 hpi, 0.5% parasitaemia) were split and treated for 4 hours with 100 nM rapamycin or DMSO (solvent control), respectively. At 20-26 hpi, the parasite populations were split, washed once in medium lacking choline chloride and AlbuMAX II and resuspended in mFA medium with or without 2 mM choline chloride to suppress or induce sexual commitment, respectively (8). After 48 hours, SCRs were quantified in the ring stage progeny based on PfAP2-G-mScarlet expression using high content microscopy. Parasites were stained with 5 µg/ml Hoechst and imaged in the DAPI and the TRITC channels to determine the proportion of sexually committed parasites (mScarlet/Hoechst double-positive cells) among all infected RBCs (Hoechst-positive cells) using the ImageXpress Micro XLS widefield high content screening system (Molecular Devices) and the MetaXpress analysis software (version 6.5.4.532, Molecular Devices) as previously described (55).

### Quantification of Pfg377 and PfARK-GFP signals in late stage gametocytes

To assess the sex-specific expression of PfARK1-GFP and PfARK2-GFP in stage V gametocytes, IFAs using mouse α-GFP and rabbit α-Pfg377 antibodies and DAPI were performed on methanol/acetone-fixed samples. Image acquisition with the 40x objective was followed by automated quantification of the fluorophore signal intensities emitted from the secondary antibodies (Alexa Fluor 488-conjugated α-mouse IgG, Alexa Fluor 568-conjugated α-rabbit IgG) and DAPI signal using a custom-built Fiji macro to score individual cells as follows. Images were segmented based on the Pfg377 signal using the auto-threshold “Triangle”, which identified potential male and female gametocytes [note that while Pfg377 is routinely used as a marker for female gametocytes, male gametocytes also stain weakly positive (60, 62)]. Regions of interest (ROIs) were defined by applying a size threshold (25-75 µm^2^) and a circularity parameter (0.35-1) to select for individual gametocytes and exclude signals derived from cellular debris/dead cells or cell aggregates. ROIs located at the image edges or associated with integrated DAPI signal intensity below an empirically defined threshold were excluded. Finally, for all remaining ROIs (gametocytes), the integrated signal intensities measured in the GFP channel (ARK1-GFP or ARK2-GFP) and the Texas Red channel (Pfg377) were plotted against each other in a scatter plot.

### Statistical analysis

The means and standard deviation of parasite multiplication rates and SCRs were calculated from three independent biological replicate experiments. Statistical significance (p-value < 0.05) was assessed using a paired two-sided Student’s t-test with the t.test function of the software R (version 4.2.0). Contour lines in Figs. 4C and 5C were plotted using the geom_density_2d function of the R package ggplot2 (81). The number of cells analyzed is specified in the respective figure legends.

## Acknowledgements

We are grateful to Pietro Alano and Alan Cowman for providing the α-Pfg377 and α-CENH3 antibodies, respectively. This work was supported by grants from the Swiss National Science Foundation to T.S.V (BSCGIO_157729, 310031_184785) and N.M.B.B. (310030_200683).

M.W., B.T.T., E.C., N.M.B.B and T.S.V. conceived and designed the experiments. M.W., B.T.T. and J.K. performed the experiments. M.W. and B.T.T. analysed the data. M.W., B.T.T., N.M.B.B. and T.S.V. wrote the manuscript. All authors contributed to the article and approved the submission.

