## Supplementary Information for "The three *Plasmodium falciparum* Aurora-related kinases display distinct temporal and spatial associations with mitotic structures in asexual blood stage parasites and gametocytes"

*contributed equally to this work

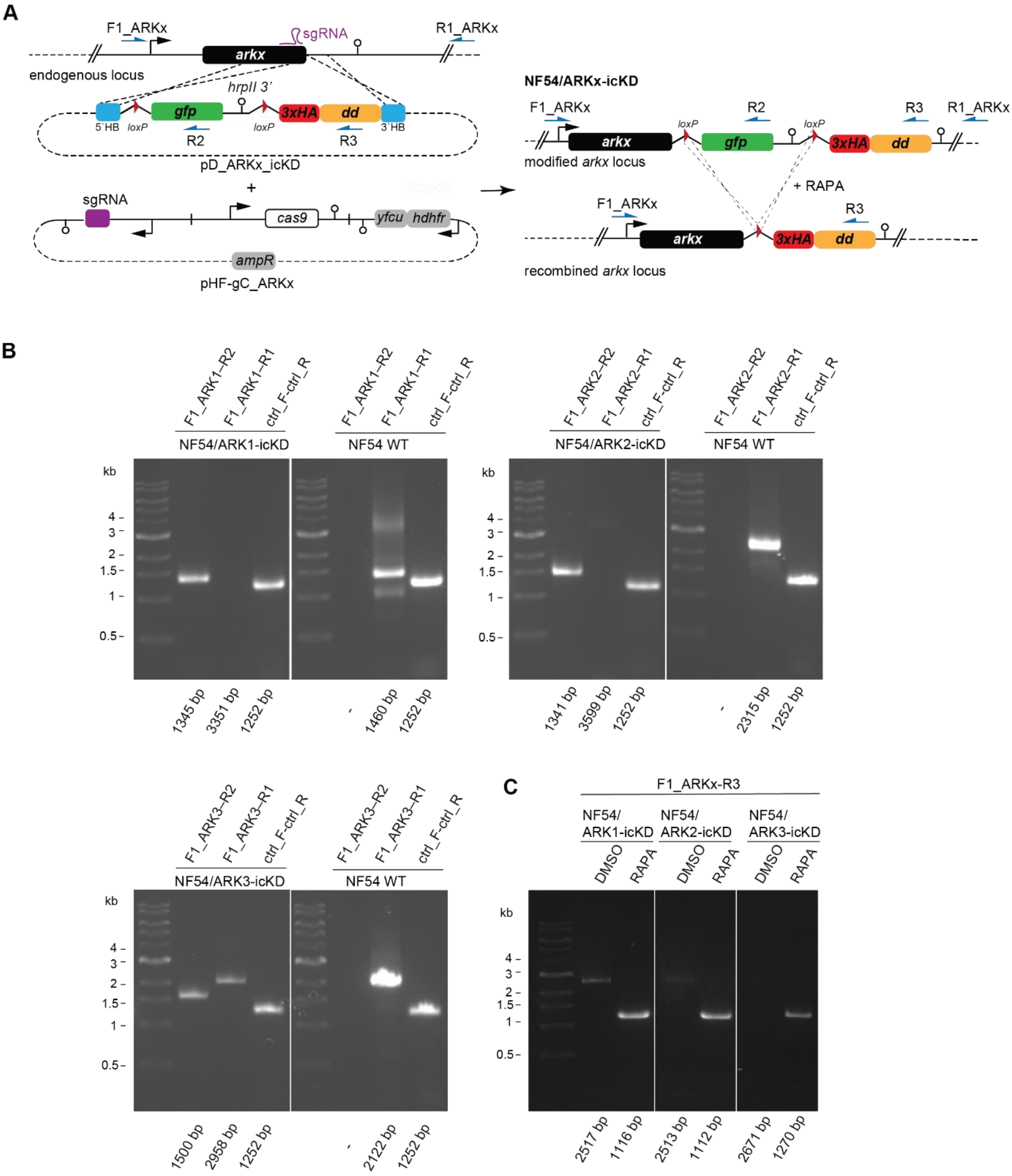


**Figure S1. CRISPR/Cas9-based engineering of PfARK1-3 icKD lines. (A)** Schematic of the *pfarkx* wild type locus in NF54/DiCre parasites and the plasmids pD_ARKx_icKD and pHF-gC_ARKx (left) that were used to generate the modified *pfarkx* loci in NF54/ARKx-icKD parasite lines (right). Addition of RAPA induces the DiCre-mediated recombination of the two *loxP* sites, thus replacing the C-terminal GFP tag with the 3xHA-DD tag. Primers used for diagnostic PCRs on gDNA are indicated. *x* represents the *pfark* variants 1-3. HB, homology box. *ark*, aurora-related kinase. *gfp*, green fluorescent protein. *3xha*, triple hemagglutinin tag. *dd*, destabilization domain. *yfcu*, yeast cytosine deaminase-uracil phosphoribosyl transferase fusion gene. *hdhfr*, human dihydrofolate reductase. *ampR*, ampicillin resistance gene. *cas9, S. pyogenes* Cas9. sgRNA, single guide RNA. RAPA, rapamycin. **(B)** Correct locus editing was confirmed by diagnostic PCRs on gDNA of NF54/ARKx-icKD parasites. Numbers at the bottom indicate the expected band sizes. **(C)** Diagnostic PCRs on gDNA of RAPA- and DMSO-treated NF54/ARKx-icKD parasites confirm successful excision of the floxed *gfp* sequence upon RAPA treatment. Numbers at the bottom indicate the expected band sizes.

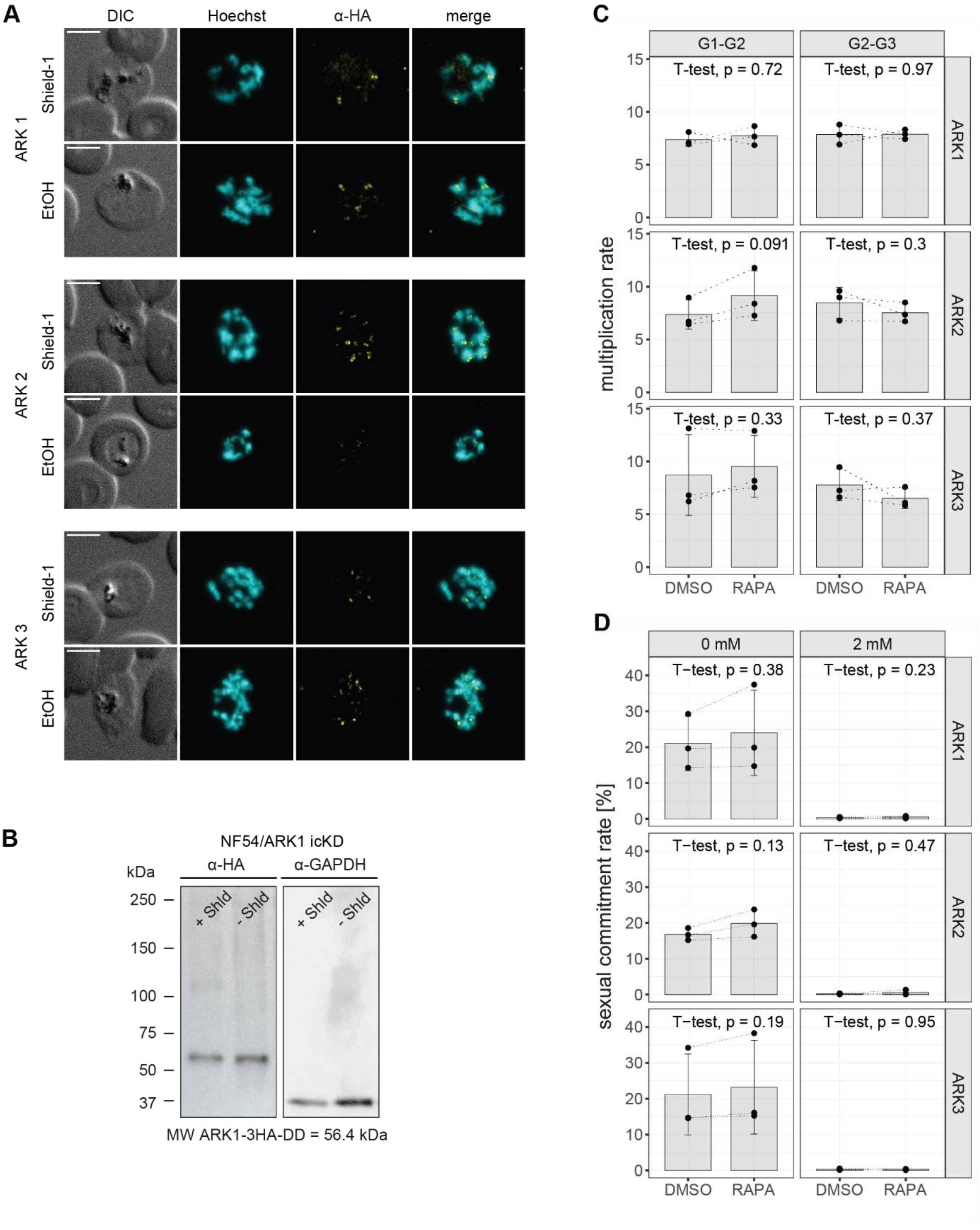


**Figure S2. Removal of Shield-1 from RAPA-treated NF54/ARKx-icKD parasites has no effect on parasite muliplication and sexual commitment rates. (A)** IFAs of RAPA-treated NF54/ARKx-icKD schizonts expressing 3xHA-DD-tagged PfARKs show that PfARK1-3xHA-DD and PfARK3-3xHA-DD expression is not noticeably reduced in the absence of Shield-1, while PfARK2-3xHA-DD expression is moderately reduced. To induce the knockdown of protein expression, parasites (0-6 hpi) were treated with RAPA for 4 hours, before the population was split and treated with Shield-1 or the vehicle control EtOH. **(B)** Expression levels of the PfARK1-3xHA-DD fusion protein were assessed by Western blot. Parasite cultures at 0-6 hpi were treated with RAPA for 4 hours, before the population was split and treated with Shield-1 or the vehicle control EtOH. Protein lysates were collected 40 hours later from late schizonts. α-GAPDH antibodies served as loading control. The expected size of the PfARK1-3xHA-DD fusion protein is 56.4 kDa. Expression of the PfARK2-3xHA-DD and PfARK3-3xHA-DD fusion proteins could not be detected by Western blot. **(C)** Parasites were split at 0-6 hpi in and treated for 4 hours with either RAPA or DMSO. For both populations, multiplication rates from generation 1 to generation 2 (G1-G2) and G2-G3 were determined by flow cytometry. Data points represent the multiplication rates from G1-G2 and G2-G3 obtained from three biological replicate experiments. Error bars represent the standard deviation. Statistical significance was assessed using a paired two-sided Student's t test. **(D)** Parasites were split at 0-6 hpi in G1 and treated for 4 hours with either RAPA or DMSO and then grown for one replication cycle in mFA in either the presence (2 mM) or absence (0 mM) of choline chloride. Sexual commitment rates were determined based on PfAP2-G-mScarlet expression in the ring stage progeny. Data points represent the proportion of PfAP2-G-mScarlet-expressing parasites among all ring stage parasites obtained from three biological replicate experiments. Error bars represent the standard deviation. Statistical significance was assessed using a paired two-sided Student's t test.

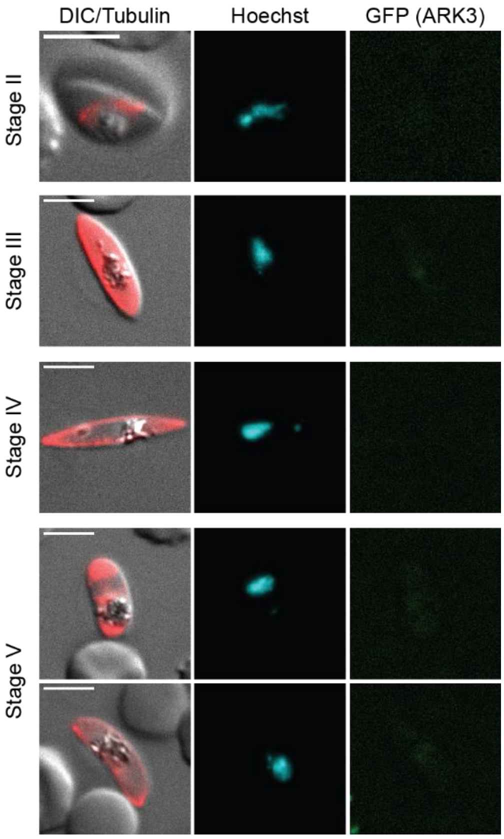


**Figure S3. PfARK3 expression was not detected in gametocytes.** PfARK3-GFP expression in gametocytes was assessed by live cell fluorescence microscopy. DNA was stained with Hoechst. MTs were stained with SPY555-tubulin. Stages of gametocyte development are indicated. Representative images are shown. DIC, differential interference contrast. Scale bar = 5 µm.

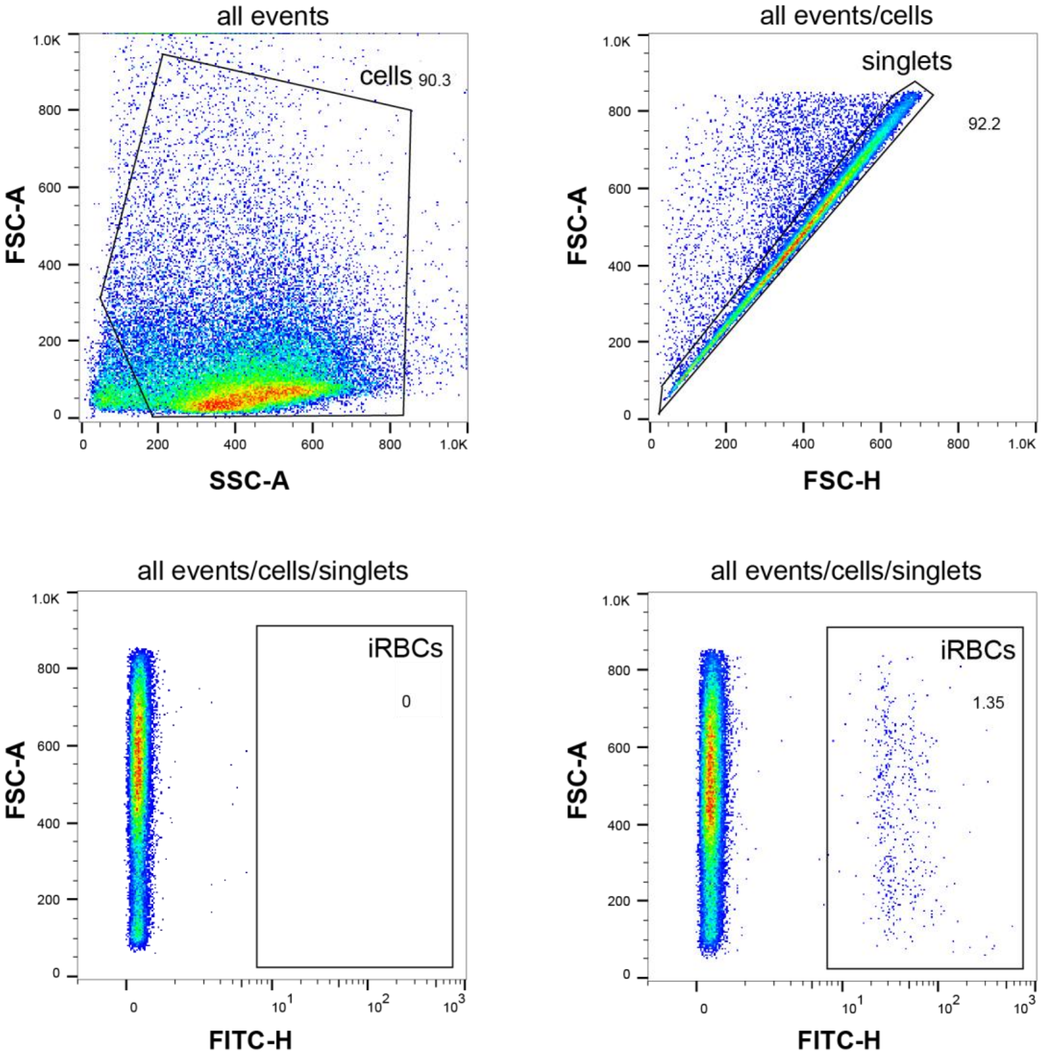


**Figure S4. Quantification of parasitaemia by flow cytometry.** The gating strategy aimed at distinguishing cells from small debris (FSC-A vs. SSC-A) and removing measurements with more than one cell per droplet (FSC-A vs. FSC-H). Infected red blood cells were identified based on their SYBR Green signal intensity (FSC-A vs. FITC-H). Representative flow cytometry data is shown. The plot at the bottom left was obtained from an unstained control, confirming the specificity of the SYBR Green signal. FSC, forward scatter. SSC, side scatter, FITC, fluorescein isothiocyanate. A, area. H, height.

**Table S1: All oligonucleotides used in this study.**

| name | sequence (5'-3') | used for | cell line/ plasmid |
|---|---|---|---|
| F1_ARK1 | ATAATATGGGCTATAAGAACATGTC | PCR on gDNA | NF54/ARK1_icKD |
| R1_ARK1 | CCAAATGGAGGGACAAAATAATG | PCR on gDNA | NF54/ARK1_icKD |
| F1_ARK2 | CGCTCTGAAGGTAATGGC | PCR on gDNA | NF54/ARK2_icKD |
| R1_ARK2 | GTTAATGCTTACATATTCCTCATC | PCR on gDNA | NF54/ARK2_icKD |
| F1_ARK3 | CAACTTTACCTAACATGAATATTGAT | PCR on gDNA | NF54/ARK3_icKD |
| R1_ARK3 | GTAGAATAAATTTGTGAAATACATAAAAG | PCR on gDNA | NF54/ARK3_icKD |
| R2 | GTGTGAGTTATAGTTGTATTCC | PCR on gDNA | NF54/ARKx_icKD |
| R3 | TCATTCCAGTTTTAGAAGCTCCAC | PCR on gDNA | NF54/ARKx_icKD |
| ctrl_F | AATGGCAGTAACAAAACTTGG | PCR on gDNA | NF54/ARKx_icKD |
| ctrl_R | CTTAGTTGTTAGTAATGTGTACGG | PCR on gDNA | NF54/ARKx_icKD |
| PCR1_F | CTGGCGTAATAGCGAAGAGG | cloning | pD_ARKx_icKD |
| PCR1_R | CATTAATGAATCGGCCAACG | cloning | pD_ARKx_icKD |
| PCR2_ARK1_F | AGCGAGTCAGTGAGCGAGGAAATAGCACATTTGGATTTGAAACC | cloning | pD_ARK1_icKD |
| PCR2_ARK1_R | TTTTTTATTTACCTATTTCCCTTATTATGGAACCAAATAGGAGTC | cloning | pD_ARK1_icKD |
| PCR3_ARK1_F | AGCTTCTAAAACTGGAATGACATATGGTTTCACAACAAATGAC | cloning | pD_ARK1_icKD |
| PCR3_ARK1_R | CCTTTTCTCTTGTGGATCCGGACATTTGTGACGATTTGTTGG | cloning | pD_ARK1_icKD |
| PCR4_ARK1_F | CTATTTGGTTCCATAATAAGGGAAATAGGTAAATAAAAAAAATAATATAC | cloning | pD_ARK1_icKD |
| PCR4_ARK1_R | ATTTGTTGTGAAACCATATGTCATTCCAGTTTTAGAAGCTCC | cloning | pD_ARK1_icKD |
| PCR2_ARK2_F | AGCGAGTCAGTGAGCGAGGAAGCTAATGGAGGATCAGTACG | cloning | pD_ARK2_icKD |
| PCR2_ARK2_R | TTTTTTATTTACCTATTTCCAATAAACTGCTTAATCCAAGGATG | cloning | pD_ARK2_icKD |
| PCR3_ARK2_F | AGCTTCTAAAACTGGAATGAAAAACAATTCATATAAAAGAAAGCG | cloning | pD_ARK2_icKD |
| PCR3_ARK2_R | CCTTTTCTCTTGTGGATCCGTATTTTCATAATCATGTTCATAGTAC | cloning | pD_ARK2_icKD |
| PCR4_ARK2_F | CTTGGATTAAGCAGTTTATTGGAAATAGGTAAATAAAAAAAATAATATAC | cloning | pD_ARK2_icKD |
| PCR4_ARK2_R | TCTTTTATATGAATTGTTTTTCATTCCAGTTTTAGAAGCTCC | cloning | pD_ARK2_icKD |
| PCR2_ARK3_F | AGCGAGTCAGTGAGCGAGGATATAAAGCAAAACAATAATACTAAGG | cloning | pD_ARK3_icKD |
| PCR2_ARK3_R | TTTTTTATTTACCTATTTCCAGTAGTTTTGTTAAGTATAGTTGCTG | cloning | pD_ARK3_icKD |
| PCR3_ARK3_F | AGCTTCTAAAACTGGAATGACAGAGAAAAATTAAATGGCAACATC | cloning | pD_ARK3_icKD |
| PCR3_ARK3_R | CCTTTTCTCTTGTGGATCCGTGTTATAATATTCTGATTCATTCCAC | cloning | pD_ARK3_icKD |
| PCR4_ARK3_F | CTATACTTAACAAAACTACTGGAAATAGGTAAATAAAAAAAATAATATAC | cloning | pD_ARK3_icKD |
| PCR4_ARK3_R | TGCCATTTAATTTTTCTCTGTCATTCCAGTTTTAGAAGCTCC | cloning | pD_ARK3_icKD |
| gRNA_ARK1_a | TATTAAACCATATGGGTGTTAGAA | cloning | pHF-gC_ARK1 |
| gRNA_ARK1_b | AAACTTCTAACACCCATATGGTTT | cloning | pHF-gC_ARK1 |
| gRNA_ARK2_a | TATTAACTGCTGAAGAAAGAATTT | cloning | pHF-gC_ARK2 |
| gRNA_ARK2_b | AAACAAATTCTTTCTTCAGCAGTT | cloning | pHF-gC_ARK2 |
| gRNA_ARK3_a | TATTATAAATAAACAGCAAAGATT | cloning | pHF-gC_ARK3 |
| gRNA_ARK3_b | AAACAATCTTTGCTGTTTATTTAT | cloning | pHF-gC_ARK3 |
